# Inheritance bias of deletion-harbouring mtDNA in yeast: the role of copy number and intracellular selection

**DOI:** 10.1101/2024.09.11.612442

**Authors:** Nataliia D. Kashko, Iuliia Karavaeva, Elena S. Glagoleva, Maria D. Logacheva, Sofya K. Garushyants, Dmitry A. Knorre

## Abstract

Eukaryotic cells contain multiple copies of mitochondrial DNA (mtDNA) molecules that replicate independently. Cell mtDNA content and variability contributes to the overall cell fitness. During sexual reproduction, fungi usually inherit mtDNA from both parents, however, the distribution of the mtDNA in the progeny can be biased toward some mtDNA variants. For example, crossing *Saccharomyces cerevisiae* strain carrying wild type (*rho*^*+*^) mtDNA with the strain carrying mutant mtDNA variant with large deletion (*rho*^−^) can produce up to 99-100% of *rho*^−^ diploid progeny. Two factors could contribute to this phenomenon. First, *rho*^−^ cells may accumulate more copies of mtDNA molecules per cell than wild-type cells, making *rho*^−^ mtDNA a prevalent mtDNA molecule in zygotes. This consequently leads to a high portion of *rho* ^−^ diploid cells in the offspring. Second, *rho*^−^ mtDNA may have a competitive advantage within heteroplasmic cells, and therefore could displace *rho*^*+*^ mtDNA in a series of generations, regardless of their initial ratio. To assess the contribution of these factors, we investigated the genotypes and phenotypes of twenty two *rho*^−^ yeast strains. We found that indeed *rho*^−^ cells have a higher mtDNA copy number per cell than *rho*^*+*^ strains. Using an *in silico* modelling of mtDNA selection and random drift in heteroplasmic yeast cells, we assessed the intracellular fitness of mutant mtDNA variants. Our model indicates that both higher copy numbers and intracellular fitness advantage of the *rho*^*-*^ mtDNA contribute to the biased inheritance of *rho*^−^ mtDNA.

## Introduction

Eukaryotic cells usually contain multiple not necessarily identical copies of mitochondrial DNA (mtDNA). This phenomenon is referred to as mitochondrial heteroplasmy. Heteroplasmy is linked to a variety of genetic diseases [1] and aging [2]. The presence of multiple mtDNA molecules within cells establishes a complex hierarchy of mtDNA populations. This hierarchy enables natural selection to operate at various levels of biological organisation, such as within individual cells and organisms, and between organisms [3–6]. As a result, “selfish” mtDNA variants may gain an advantage within individual cells, despite being neutral or exhibiting deleterious effects at the cellular or organismal level [5,7–9].

Several experimental observations support the idea that mtDNA selection acts on different levels of organisation. First, pathogenic mitotypes containing mutations in the D-loop, a non-coding region of mtDNA controlling the initiation of its replication [10], gradually replace donor mtDNA during the development of a multicellular organism from an embryo [11]. Second, the common starling population in Australia was shown to carry two coexisting mtDNA variants (mitotypes). One of the mitotypes demonstrated a moderate advantage at the intracellular level while decreasing the fitness of the whole organism [12]. Next, natural isolates of nematodes *Caenorhabditis briggsae* harbour two distinct mtDNAs: a full-length variant and a deleterious *Δnad5* variant lacking a portion of the *NAD5* gene. The proportion of *Δnad5* mtDNA increased across generations if the populations underwent frequent bottlenecks [13]. Finally, it has been shown that in the baker’s yeast *Saccharomyces cerevisiae*, the selection of mtDNA also depends on population size. In an experimental evolution study, a mutant deleterious mtDNA variant displaced the parental wild-type variant in small populations, but not in the larger ones [14].

*S. cerevisiae* has been proven to be an invaluable model for exploring intracellular mtDNA selection. This yeast species can proliferate under fermentative conditions in the presence of mtDNA mutations that disrupt oxidative phosphorylation (referred to as the *rho*^−^ genotype) or in the complete absence of mtDNA (*rho*^*0*^ genotype). Laboratory yeast strains often produce *rho*^−^ mutants, i.e. having *rho*^−^ genotype of their mtDNA, which usually contain large deletions in their mtDNA sequences [15,16]. At the same time, yeast cells with *rho*^−^ and *rho*^*0*^ genotypes can be differentiated from the wild type (*rho*^*+*^) cells by their *‘petite’* phenotype. This phenotype is defined by slow growth and an inability to utilise non-fermentable carbon sources [17]. Additionally, *S. cerevisiae* has biparental mtDNA inheritance, where both gametes (mating haploid cells) transmit their mtDNAs to the diploid progeny [18], which allows easy production of heteroplasmic yeast cells.

Intriguingly, in the case of some *rho*^−^ mitotypes, the *rho*^−^ X *rho*^*+*^ cell crossings produce mainly *petite* diploid cells [19]. This phenomenon, known as mtDNA suppressivity, suggests that *rho*^−^ mtDNAs might be positively selected, even though it would cause a detrimental (*petite*) phenotype. However, the relative contribution of these factors to mtDNA suppressivity remains not fully characterised. On one side, *rho* ^−^ mtDNAs could have a replication rate advantage in heteroplasmic cells. Indeed, some *rho*^−^ mtDNAs incorporate radioactively labelled nucleotides faster than the *rho*^*+*^ full-length molecules [20]. On the other side, *rho*^−^ haploid strains typically contain a significantly higher quantity of mtDNA molecules per cell [21]. The increased number of mtDNA molecules in *rho*^−^ cells compared to *rho*^*+*^ cells may explain the preferential inheritance of *rho*^−^ mitotypes in *rho*^−^ *x rho*^*+*^ crossings, even in the absence of intracellular selection.

In this study, we generated a set of 22 *rho*^−^ *S*.*cerevisiae* strains, mapped deletions in their mtDNA, analysed mtDNA copy number per cell, and suppressivity for each mutant. To assess mtDNA copy number contribution to the degree of suppressivity, we simulated mtDNA selection and genetic drift during crossings using a stochastic model. Utilising this model, which incorporates estimates of mtDNA copy numbers per cell and growth rates of petite cells, we calculated possible suppressivity values under the assumptions of neutral drift and varying levels of replication advantage for *rho*^−^ mtDNAs. Together, our experimental data and simulations support the assumption that intracellular selection favours *rho*^−^ mtDNAs in heteroplasmic yeast cells.

## Results

### mtDNA copy number in rho− strains correlates with their suppressivity

To identify the genomic features that contribute to suppressivity, we generated 23 spontaneous *rho*^−^ strains from a parental haploid strain [mat alpha *rho*^*+*^ *URA*^*+*^ *leu*^−^]. To measure their suppressivity we crossed those strains with the *rho*^*+*^ strain of the opposing mating type [mat a *rho*^*+*^ *ura*^−^ *LEU*^*+*^]. We defined the suppressivity of the *rho*^−^ strains as the percentage of diploid progeny cells that are unable to grow on glycerol (Figure 1A). One of the low-suppressive strains was found to be *rho*^*0*^ and was therefore excluded from further analysis. The suppressivity of the remaining strains ranged from 19.4 to 90.2 percent (Table S1).

**Figure 1.**
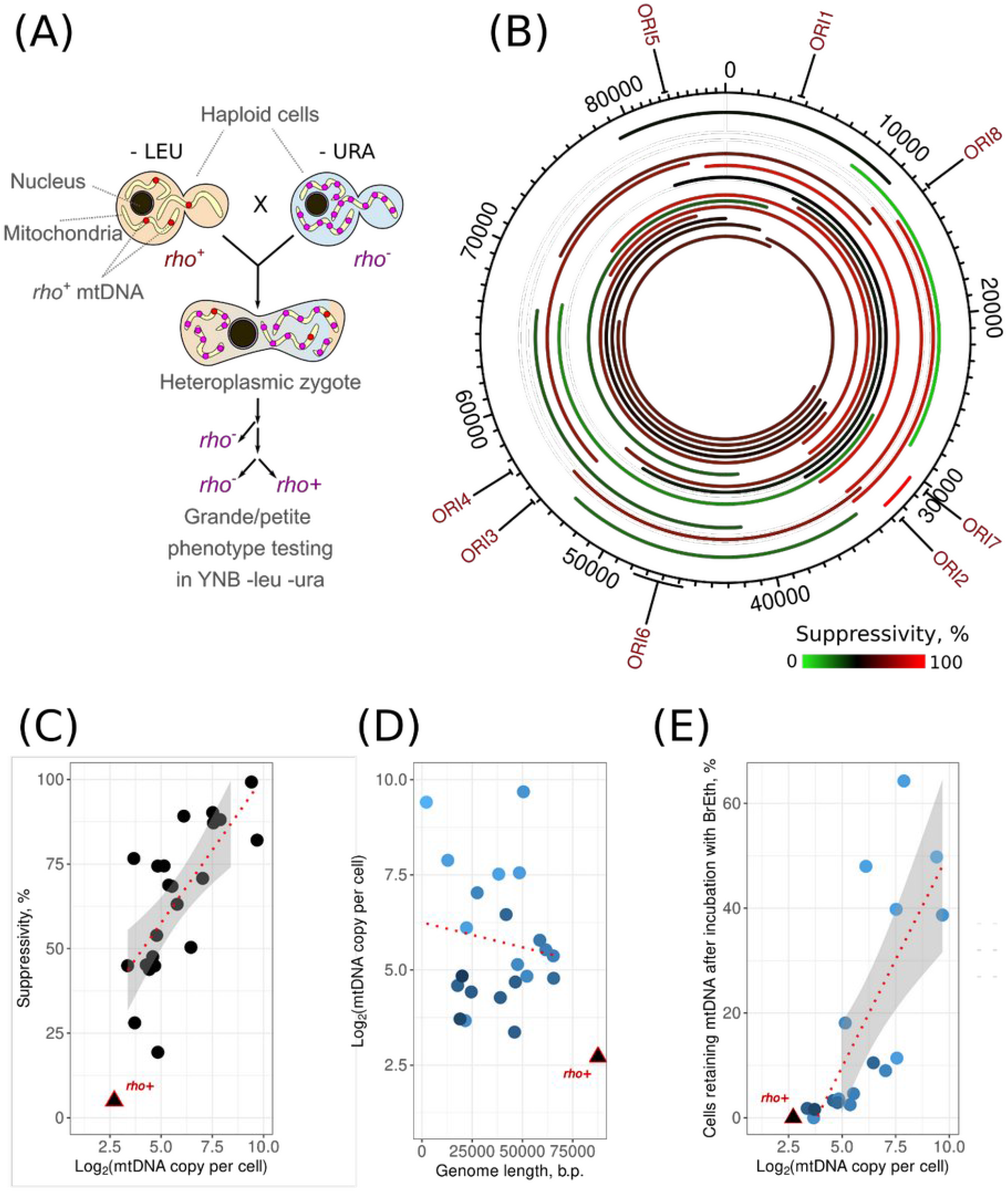
Suppressivity of spontaneous *rho*^−^ strains. **(A)** Experimental design for estimating suppressivity; **(B)** *S. cerevisiae* mitochondrial DNA map featuring the locations of replication origins. The arcs on the map illustrate the mtDNA segments preserved in various spontaneous *rho*^−^ mutants examined in this study. The colour of the arcs corresponds to the average suppressivity of the respective *rho*^−^ strains. **(C)** Correlation of *rho*^−^ mtDNA copy number per nuclear genome estimated from NGS data with suppressivity (Kendall’s rank correlation tau = 0.471, p-value = 0.001); **(D)** *rho*^−^ mtDNA genome size does not correlate with mtDNA copy number per nuclear genome; **(E)** The ability of *rho*^−^ yeast cell to retain mtDNA upon growth with the DNA-intercalating agent ethidium bromide correlates with *rho*^−^ mtDNA copy number (Kendall’s rank correlation tau = 0.72, p-value = 2*10^-5^). Data for the *rho*^*+*^ strain is shown in all plots but is not included in the linear model and correlation tests.

Next, we isolated total DNA from the *rho*^−^ strains and performed whole-genome sequencing to identify the position of deletions in their mtDNA. We show that obtained *rho*^−^ mtDNA mutants vary in the deletion length and position (Figure 1B). In 7 out of 22 strains the position of the primary deletion was confirmed by Sanger sequencing (Table S1). We found that the studied strains retained 76.8 to 14.8 percent of the unique parental mtDNA sequence. All strains retained at least one of the eight predicted replication origins (Figure 1B). We also sequenced the previously isolated hypersuppressive (HS) strain which showed on average 99% suppressivity and retained only a short 2.2 kb fragment of mtDNA spawning a region flanked by *ORI2* and *ORI7* [22]. In our set of strains, we did not find a strong association between the retained replication origins and suppressivity, however, it may be noticed that two low-suppressive strains, namely *IIa10* and *13*, lack all three active replication origins: *ORI2*, *ORI3*, and *ORI5* (Figure 1B).

To assess the relative mtDNA copy number for each *rho*^−^ strain, we compared the average read coverage depth for mtDNA with the one for the nuclear genome (nDNA, Table S2). However, due to the AT-rich nature of the yeast mitochondrial genome, the mapping algorithms artificially map less reads to AT-rich regions that lead to non-uniform read coverage along mtDNA (Figure S1). To overcome this issue, other studies routinely assess mtDNA copy number only in the region of high GC-content spanning from position 14,000 to 20,000 [23], and where indeed read depth is uniform (Figure S1). Unfortunately, this region was absent in some of our *rho*^−^ strains. Thus, we selected three short mtDNA regions (coordinates 8002-8153, 31222-31305, 48195-48296, based on the reference yeast mitochondrial genome) to ensure that each *rho*^−^ strain retains at least one of these regions (Table S2). To estimate the mitochondrial DNA copy number in our *rho*^−^ strains, we estimated the copy number for each of the available regions and selected the highest value among them. We validated results obtained from the procedure described above for a subset of *rho*^−^ strains by quantitative PCR with three pairs of primers designed for the regions used for the NGS copy number assessment (Table S3, Figure S1). NGS and qPCR approaches for quantification of mtDNA/nDNA ratios showed concordant results, the values were positively correlated with Kendall’s rank correlation tau = 0.683, p-value = 8.2×10^-5^ (Figure S2). However, the absolute values of the ratios mtDNA/nDNA obtained with NGS and qPCR methods substantially differed (see Table S2 and Table S4). For instance, according to NGS, the mtDNA copy number of the control wild-type strain ranged from an unrealistic 3.0 to 15.5 copies per haploid nuclear genome, depending on the mtDNA region selected as reference (Figure S2). In contrast, qPCR provided a higher estimate of 22.2-26.6 mtDNA copies per nuclear genome (Table S4)

We used the NGS-based evaluation of the mtDNA copy number and correlated it with the corresponding strain’s suppressivity. We showed that mtDNA/nDNA ratios of the *rho*^−^ strains positively correlate with their average suppressivity (Figure 1Cl; Kendall’s rank correlation tau = 0.47, p-value = 0.001). At the same time, we found no correlation between mtDNA copy number and the length of the mtDNA remaining in the cells (Figure 1D). Since both methods for determining the mtDNA/nDNA ratio (qPCR-based and NGS-based) relied on DNA amplification, we decided to complement those measurements by another approach. We hypothesised that *rho*^−^ str ains with increased mtDNA copy number would be more resilient to the loss of mtDNA in response to the inhibition of mtDNA replication. To test this, we treated *rho*^−^ strains with the DNA-intercalating agent, ethidium bromide. Then, we stained the cells with DAPI to visualise nuclear and mitochondrial DNA in the cells. Our findings confirmed that *rho*^−^ cells with high mtDNA/nDNA ratios were more resistant to mtDNA loss (Figure 1E). In the analysed set of *rho*^−^ strains, the percentage of *rho*^−^ cells retaining mtDNA after incubation with the ethidium bromide correlated with the average mtDNA/nDNA ratios in the strain (Figure 1E, Kendall’s rank correlation tau = 0.72, p-value = 2*10^-5^) and suppressivity (Kendall’s rank correlation tau = 0.56, p-value = 8*10 ^-4^, Figure S3). Taken together, three independent approaches to estimate mtDNA copy number show a positive correlation between the mtDNA copy number and suppressivity.

### Suppressivity drifts after genetic bottleneck

To explore the effects of genetic drift and selection on *rho*^−^ mtDNA variants in yeast cell populations, we tested whether suppressivity remains constant in *rho*^−^ strains or is subject to drift. To test this, we took several strains with varying suppressivity from our set and derived subclones by selecting colonies that presumably originated from a single cell. We evaluated suppressivity of these subclones as described earlier. Figure 2 shows that the suppressivity of a parental strain that did not undergo a bottleneck by a subclone selection remained stable in independent experiments. At the same time, some subclones obtained from single cells showed convergent suppressivity which, however, strongly differed from the average suppressivity of the parental *rho*^−^ strain. This suggests the existence of secondary mtDNA mutations in *rho*^−^ strains that affect their suppressivity.

**Figure 2.**
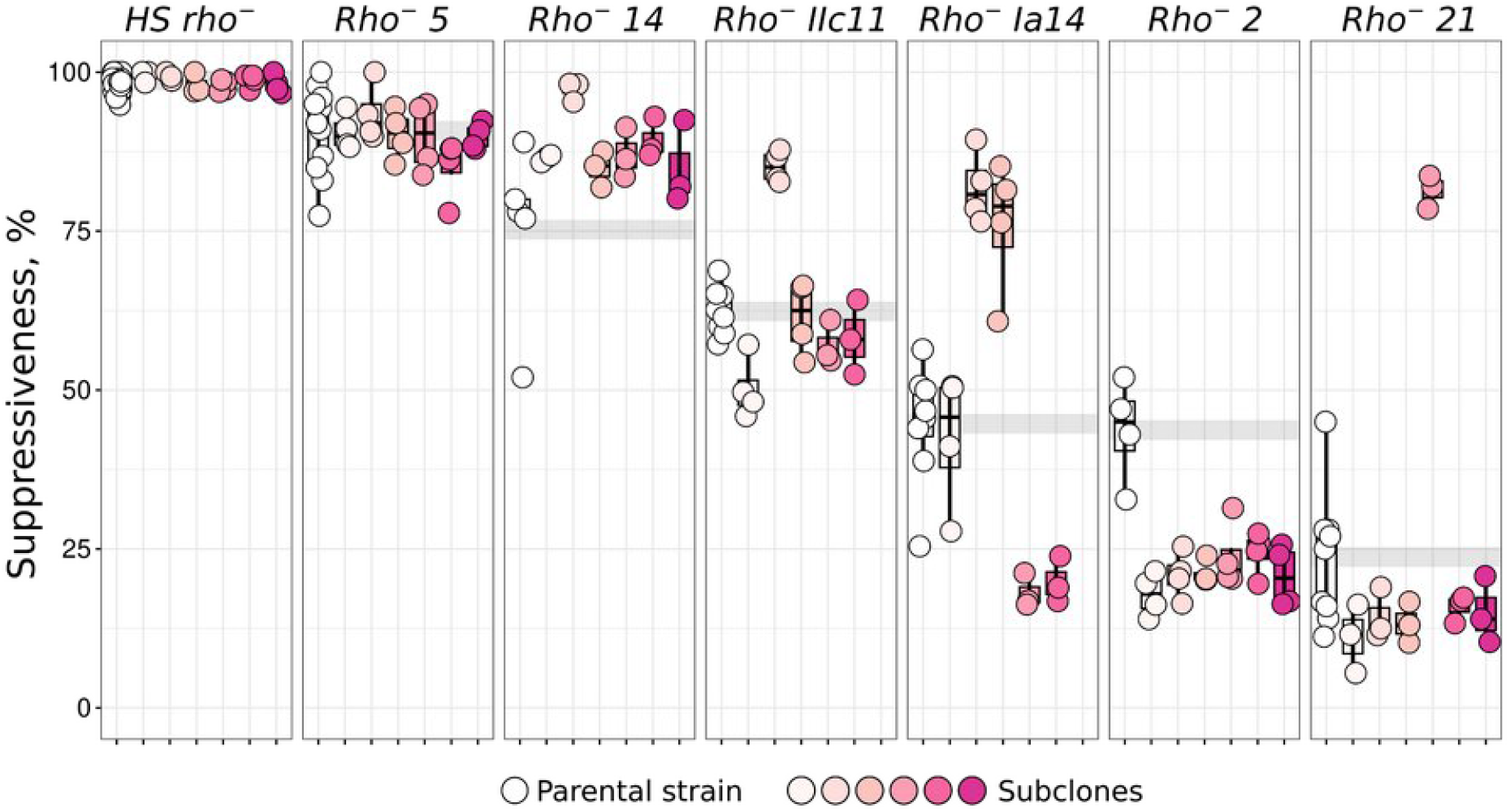
Genetic bottlenecks change the suppressivity. Suppressivity was measured in parental *rho*^−^ yeast strains and their subclones derived from each strain after a narrow genetic bottleneck (single colonies).

Our experiments demonstrated that *rho*^−^ strains can alter their suppressivity following a genetic bottleneck, prompting us to investigate whether there is a consistent trend towards increased or decreased suppressivity. To examine this, we performed a post hoc analysis of the suppressivity of 76 strains previously obtained in our laboratory. The initial assessment was conducted immediately after the narrow bottleneck, following the isolation of each *rho*^−^ strain (Figure S4). Subsequent analyses were carried out after an additional (non controlled) number of generations. We found that the initial suppressivity assessment was on average lower (5.9%±10.9) than the subsequent ones (Figure S4). Wilcoxon signed-rank test with continuity correction rejected the hypothesis that the medians of first and subsequent assessments are equal (P-value = 9.9×10^-6^). At the same time, suppressivity increased to a higher extent in the strains with low initial suppressivity (Figure S4).

### rho^−^ mtDNA suppressivity does not affect the strain growth rate

In our experimental setup for the suppressivity assessment, variation in *rho*^−^ strain suppressivity could be influenced by the difference in growth rates. For example, if a highly suppressive *rho*^−^ strain has a higher proliferation rate than a low suppressivity *rho*^−^ strain, diploid strains with homoplasmy of the highly suppressive *rho*^−^ variant can undergo more divisions than the corresponding low suppressivity *rho* ^−^ strain. As a result, it would produce more *petite* cells in the suppressivity assay. To test this possibility, we measured the maximal growth rate of our *rho*^−^ and *rho*^*+*^ strains, as well as *rho*^*0*^ cells as a control. We found no pronounced differences in growth rate between *rho*^−^ and *rho*^*0*^ strains, while the *rho*^*+*^ strain expectedly displayed higher maximal growth rate (Figure 3A, B). As the conditions during crossings deviate from those in an exponentially growing yeast suspension, we additionally tested the change in the number of *URA*^*+*^ cells (colony-forming units) before and after the crossing experiments (Figure 3C). Surprisingly, there was no difference in growth rates of *rho*^*+*^ and *rho*^−^/*rho*^*0*^ strains under these conditions, as the number of *URA*^*+*^ cells increased approximately tenfold regardless of mtDNA type for all tested strains/crossings (Figure 3D).

**Figure 3.**
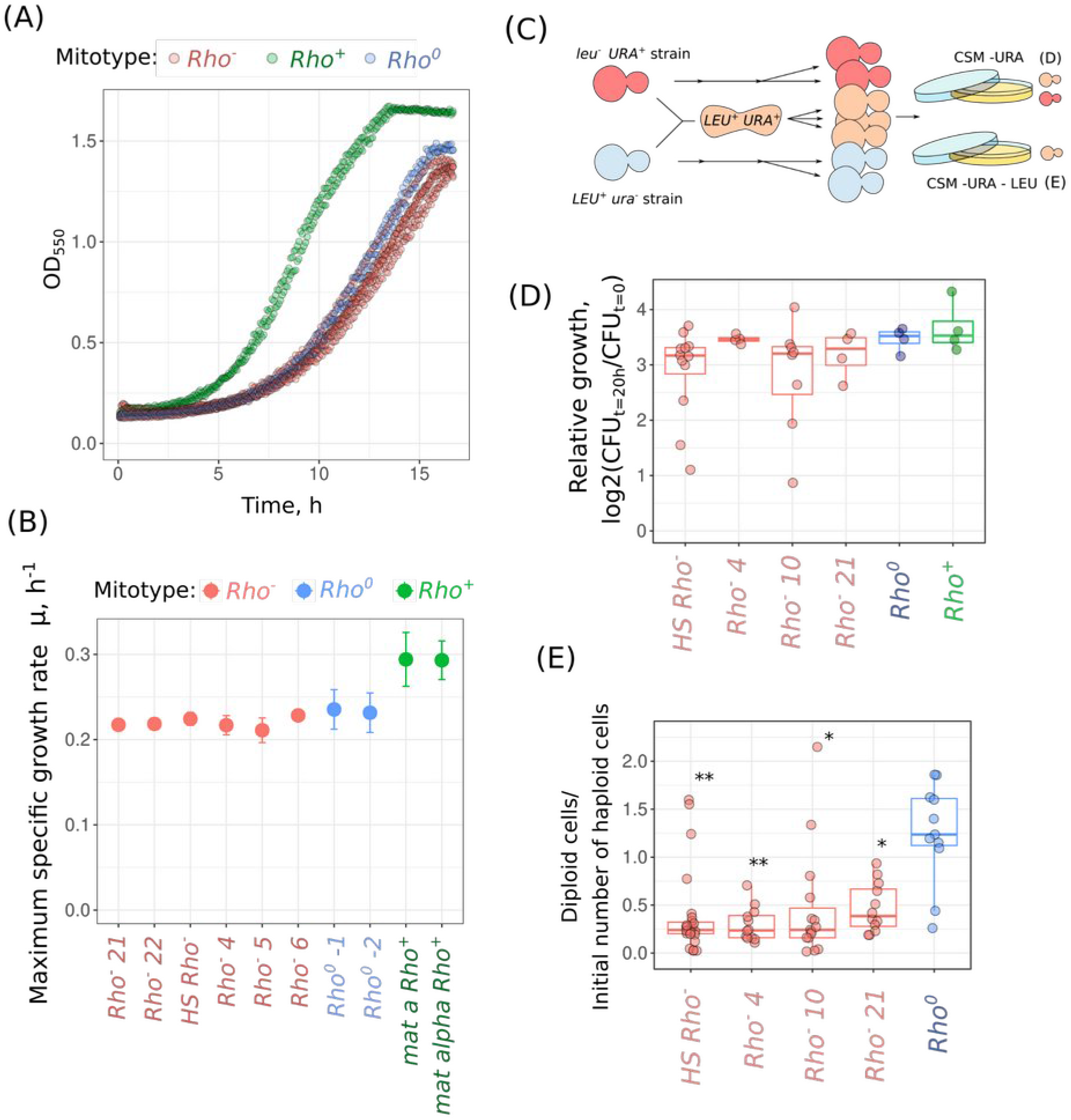
Phenotypes of *rho*^*+*^, *rho*^−^ and *rho*^*0*^ strains during the suppressivity assay experiments. Representative growth rate **(A)** and maximum specific growth rates **(B)** of *rho*^*+*^, *rho*^−^ and *rho*^*0*^ strains **(C)** Experimental scheme to assess strains growth rate during crossings; **(D)** Relative growth rates of yeast strains specified on the plot (*URA*^*+*^ *leu*^−^) c rossed with *rho*^*+*^ (*ura*^−^ *LEU*^*+*^) strain; **(E)** *URA*^*+*^ *LEU*^*+*^ diploid cell count produced by crossing indicated strain with wild-type *rho*^*+*^ strain, normalised to initial cell count; *P < 0.01, **P < 0.001 (Wilcoxon rank sum exact test) comparing *rho*^−^ X *rho*^*+*^ crosses to the *rho*^*0*^ X *rho*^*+*^.

We reasoned that the similar growth rates observed between *rho*^*+*^ and *rho*^−^/*rho* ^*0*^ strains under crossing conditions stem from the inhibition of *rho*^*+*^ cell growth by alpha factor, which is secreted by cells of the opposite mating type. In this case, the presence of functional mitochondria in *rho*^*+*^ cells might be a non-limiting factor for the growth rate. Hence, we traced the number of diploid cells during the crossing experiment. We calculated the proportion of diploid cells divided by the number of haploid cells at the beginning of the experiment (Figure 3E). We found that the growth rate of diploids obtained with *rho*^*+*^ X *rho*^*0*^ crossings exceeded that from the *rho*^*+*^ X *rho*^−^ crossings. This result shows that the presence of *rho*^−^ mtDNA in the diploid cells inhibits their growth.

### Stochastic model suggests an intracellular fitness advantage of rho^−^ mtDNAs with deletions

Our experiments yielded estimates for three parameters of *rho*^−^ cells: suppressivity, their mtDNA copy number per cell, and growth rates in crossing experiments. We reasoned that a strain’s suppressivity could be primarily derived from its growth rate and mtDNA copy number. However, it can also be influenced by the intracellular fitness of the *rho*^−^ mtDNA variant if this fitness value deviates significantly from one. In order to explore the association between these parameters, we developed a stochastic model that simulates the genetic drift and selection of mtDNAs in the heteroplasmic cells. This model takes as an input the following parameters: (1) *relative growth rates* of *rho*^*+*^ and *rho*^−^ strains; (2) the initial proportion of two mtDNA variants ( *heteroplasmy level*), with the range of possible values taken from mtDNA copy number estimations; (3) *intracellular fitness* (imputed from the value in a range from 0 to 3). In our model, intracellular fitness was defined as the relative increase in frequency of the *rho*^−^ mtDNA variant compared to the *rho*^*+*^ mtDNA variant within a single heteroplasmic cell during one cell division; The model simulates random drift and selection on intracellular and intercellular levels and returns the distribution of mtDNAs in the population of cells following several cellular divisions (see Figure S5 for the in detail model description). At the end of the simulation, the model yields *suppressivity* as the ratio of cells devoid of wild type mtDNA to the total number of cells. We ran these simulations to estimate the suppressivity with varying starting parameters of the initial heteroplasmy level and intracellular fitness of *rho*^−^ mtDNA. Figure 4A illustrates the relationship between suppressivity and input parameters: intracellular fitness and heteroplasmy levels. Given that intracellular fitness was an unknown variable in our simulations, we replotted the data using alternative coordinates (Figure 4B). In this representation, suppressivity is plotted against the heteroplasmy level, with the model-imputed intracellular fitness shown by a colour gradient. We subsequently overlaid our experimental data points for all 22 *rho*^−^ strains onto this plot.

**Figure 4.**
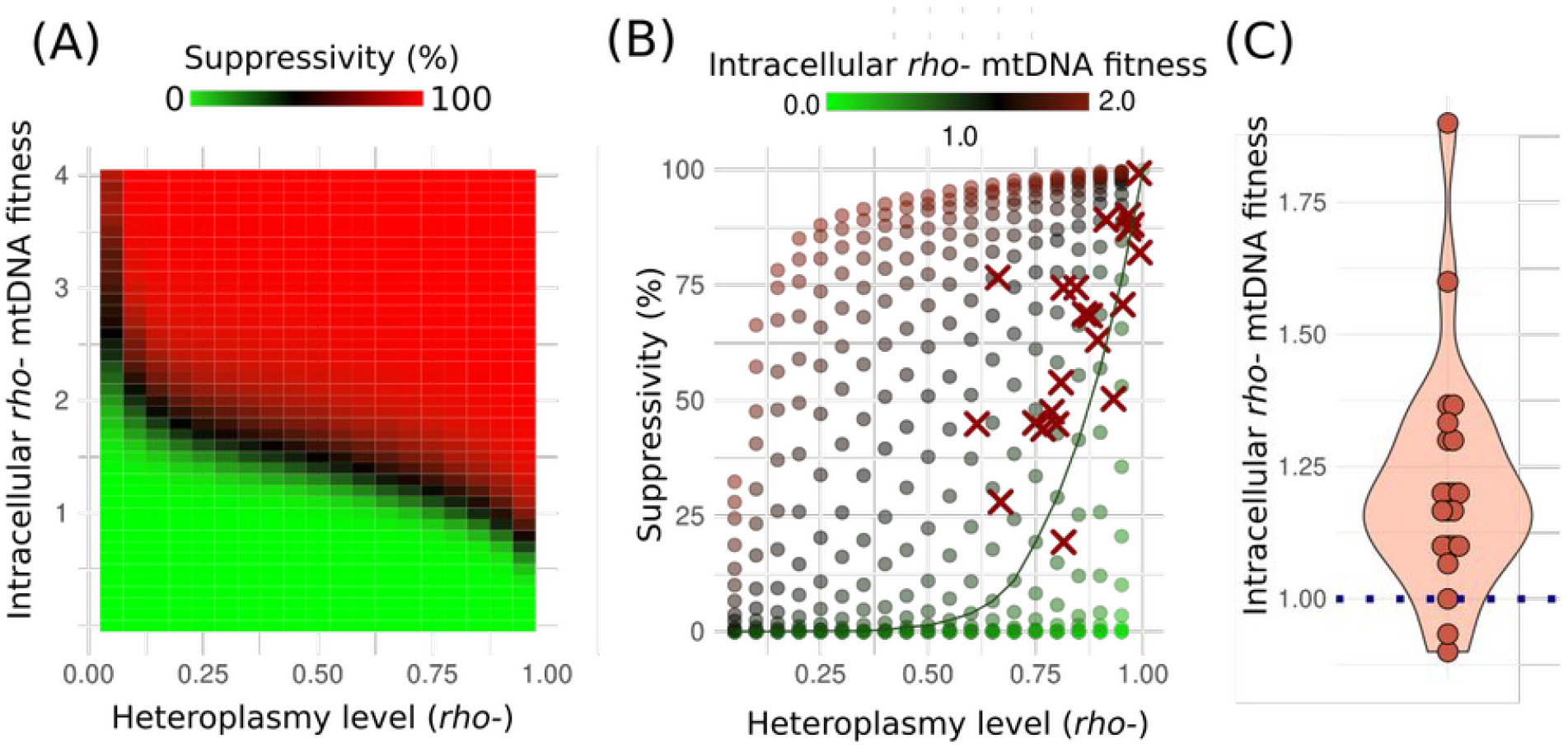
Stochastic modelling suggests that the starting proportion of *rho*^−^ and *rho*^*+*^ mtDNA molecules in the zygotes cannot explain the observed suppressivity values. **(A)** *in silico* simulation provides suppressivity estimate for varying intracellular fitness and starting heteroplasmy level parameters; **(B)** Simulation data and experimental data plotted in the coordinates of starting heteroplasmy level ∼ suppressivity; Points represent simulated data, crosses represent experimental data points for the 22 *rho*^−^ strains. A line connects the points obtained with the simulation with no replication advantage of *rho*^−^ mtDNA (Intracellular *rho*^−^ mtDNA fitness equal to 1.0); **(C)** Relative intracellular fitness of *rho*^−^ mtDNA variants calculated from the k-nearest neighbours in (B). P = 0.00006 for the hypothesis that the population distribution mean is equal to 1.0 according to Wilcoxon signed rank test with continuity correction.

Figure 4B shows that most of our experimental data points fall above the green line representing a neutral assumption that the intracellular fitness values of the *rho*^−^ and *rho*^*+*^ mtDNAs are equal, indicating that most *rho*^*-*^ mtDNAs have an intracellular fitness advantage compared to parental *rho*^*+*^ mtDNA. Furthermore, we estimated the intracellular fitness levels for our set of 22 *rho*^−^ strains from the simulation data. To do this, we determined k-nearest neighbours for each strain in the coordinates (heteroplasmy level ∼ suppressivity) (k = 6) and took their average intracellular *rho*^−^ mtDNA fitness as the estimate for our strains. Figure 4C shows the distribution of the calculated fitness values and suggests that on average, the value is significantly higher than one (fitness equal to one is the neutral expectation, assuming that there is no intracellular fitness advantage of *rho*^−^ mtDNA variants).

In our model, we incorporated several assumptions. First, we posited that mutant *rho*^−^ mtDNAs have a threshold effect on yeast growth rate. Specifically, we proposed that yeast cells switch from slow to high growth when the proportion of *rho*^*+*^ mtDNA exceeds 0.5. This assumption aligns with the generally accepted notion that deleterious effects of mtDNAs, including *rho-*, are manifested when the proportion of mutants is above a certain threshold [24]. Second, we imputed into the model the copy number per cell equal to 20 which was concordant with our estimates (Figure 1) and the published data [25]. To test the robustness of our estimates, we varied pathogenicity thresholds for *rho*^−^ mtDNAs (Figure S6) and mtDNA copy number per cell at the moment of cell division (Figure S7) in the simulations. Using the same approach mentioned above, we calculated the intracellular fitness advantage of *rho*^−^ mtDNA variants using simulations with different mtDNA copy numbers. The mean value of *rho*^−^ mtDNA intracellular fitness remained above one (Figure S8), suggesting that there is an advantage within the cell for mutant mtDNAs compared to parental wild-type mtDNA and this result is robust. Overall, our experiments and the model show that on average spontaneous deletion in yeast mtDNA provides it with a selection advantage within the cell.

## Discussion

Uniparental inheritance of mtDNA in most metazoan species prevents mtDNA recombination and imposes a problem of deleterious mutation accumulation in the mitogenomes [26]. Severe mtDNA copy number bottlenecks in the germline slow down this process [27]. In contrast to metazoa, fungi usually inherit mtDNA from both parents [28,29]. In addition, fungal mtDNA experience frequent recombination in experimental conditions [30] and natural yeast populations [31]. Biparental inheritance enables recombination which is likely to diminish the problem of mutation accumulation [27]. However, this makes the fungi prone to selfish mitochondrial genetic elements, which show preferential transmission to the progeny despite their deleterious effects on the whole organism. The simplest premise for the transmission bias of mtDNA variants is an increase in its copy number in one of the haploid cells—gametes. This would resemble the anisogamy typical for metazoa, where female gametes contain more mitochondria and mtDNA compared to male gametes [32].

To unravel how the mtDNA copy number in wild-type and *rho*^***−***^ *S. cerevisiae* cells affects their inheritance bias, we studied several *rho*^***−***^ strains harbouring large deletions in their mitochondrial genomes. mtDNA copy number per cell in all studied *rho*^−^ haploid strains exceeded that of the parental *rho*^*+*^ strain (Figure 1C). This observation is concordant with a previous study showing that *rho*^−^ yeast cells accumulated mtDNA to a copy number ∼80 times higher than that in a wild-type strain [21]. We speculate that a negative feedback loop mechanism induces mtDNA biogenesis to compensate for a deficiency in mitochondrial function that eventually leads to excessive accumulation of mtDNA. A similar mechanism was described in nematodes, where the mitochondrial unfolded protein response transcription factor ATFS-1 helped to maintain high levels of mtDNA variant harbouring a deletion by enhancing the binding of mtDNA polymerase to mutated mtDNA molecules [33]. To the best of our knowledge, such a mechanism is still unknown in yeast. Meanwhile, deletion of *COX5* gene, encoding an essential component of cytochrome oxidase, the terminal part of the respiratory chain, increases mtDNA copy number per cell [34]. Furthermore, we have recently shown that *rho*^−^ cells are usually more heterogeneous by the level of mtDNA than their parental *rho*^*+*^ cells [35]. Together, these observations suggest that the ability to encode all essential components of the OxPhos is necessary for the maintenance of the constant mtDNA levels in cells across the generations.

The simplest explanation of the suppressivity phenomena could be the observed high copy number of mtDNA molecules in the *rho*^−^ strains followed by random segregation of the molecules (Figure 1C). However, by imputing mtDNA/nDNA ratios and relative growth rates of *rho*^*+*^ and *rho*^−^ strains we were able to explain the *rho*^−^ strains’ variance in suppressivity only partially. It should be mentioned that, in yeast mtDNA molecules are usually concatenated [36,37]. Meanwhile, both methods (qPCR and NGS) used in our study to assess mtDNA copy numbers cannot distinguish molecules containing one repeat unit from concatenated molecules. To estimate the intracellular fitness of *rho*^−^ mtDNA variants, we didn’t take into account that *rho*^−^ mtDNAs consist of a smaller number of segregating units. However, if this were the case, we would overestimate the ratio of *rho*^−^ to *rho*^*+*^ mtDNA in zygotes (the heteroplasmy parameter of the model) and underestimate intracellular fitness in our prediction. In other words, even if each *rho*^−^ mitotype was represented by a separate molecule segregating independently during cell division, we could not explain the observed values of suppressivity by the initial ratio of mtDNA molecules in *rho*^−^ and *rho*^*+*^ strains crossing. Therefore, we conclude that in yeast, some mtDNA variants with large deletions can have pronouncedly higher intracellular fitness than full-size wild type mtDNAs.

In line with the conclusion that some *rho*^−^ mtDNAs can have higher intracellular fitness than others, we have shown that the suppressivity of yeast *rho*^−^ clones can be changed after the genetic bottleneck (Figure 2) and, on average, increase with generations (Figure S4). This increase can be attributed to two factors. First, cells with secondary mutations in mtDNA with high suppressivity proliferate faster than parental cells with original mutation. However, Figure 3 shows that there is no pronounced difference between the growth rate of *rho*^−^ strains with different deletions. Second, spontaneous secondary deletions in *rho*^−^ strains could lead to the appearance of mtDNA variants with intracellular fitness exceeding the intracellular fitness of the original *rho*^−^ mitotype. We suggest that the second scenario is more likely than the first one. Indeed, considering that different *rho*^−^.mtDNA molecules can have various intracellular fitness, e.g. due to the difference in replication origin densities, eventually *rho*^−^ cells will produce new *rho*^−^ variants with increased intracellular fitness. Due to an increase in intracellular fitness, this variant is expected to displace parental *rho*^−^ variants within the same cell. Meanwhile, given that there is no exchange of mtDNA between cells (while they are not crossing) and no selection between *rho*^−^ cells at the whole cell level, this variant has a chance to displace the parental variant in all lineages due to the genetic drift. As a result, newly emerging mtDNA variants with low intracellular fitness are expected to be outcompeted and removed from the population, while the mtDNA with high intracellular fitness will displace the parental variants (Figure 5). We propose that as a result, a ratchet-like mechanism leads to a gradual increase of cells carrying suppressive *rho*^−^ mtDNA in small yeast populations to a certain limit based on a maximal possible replication origin density.

**Figure 5.**
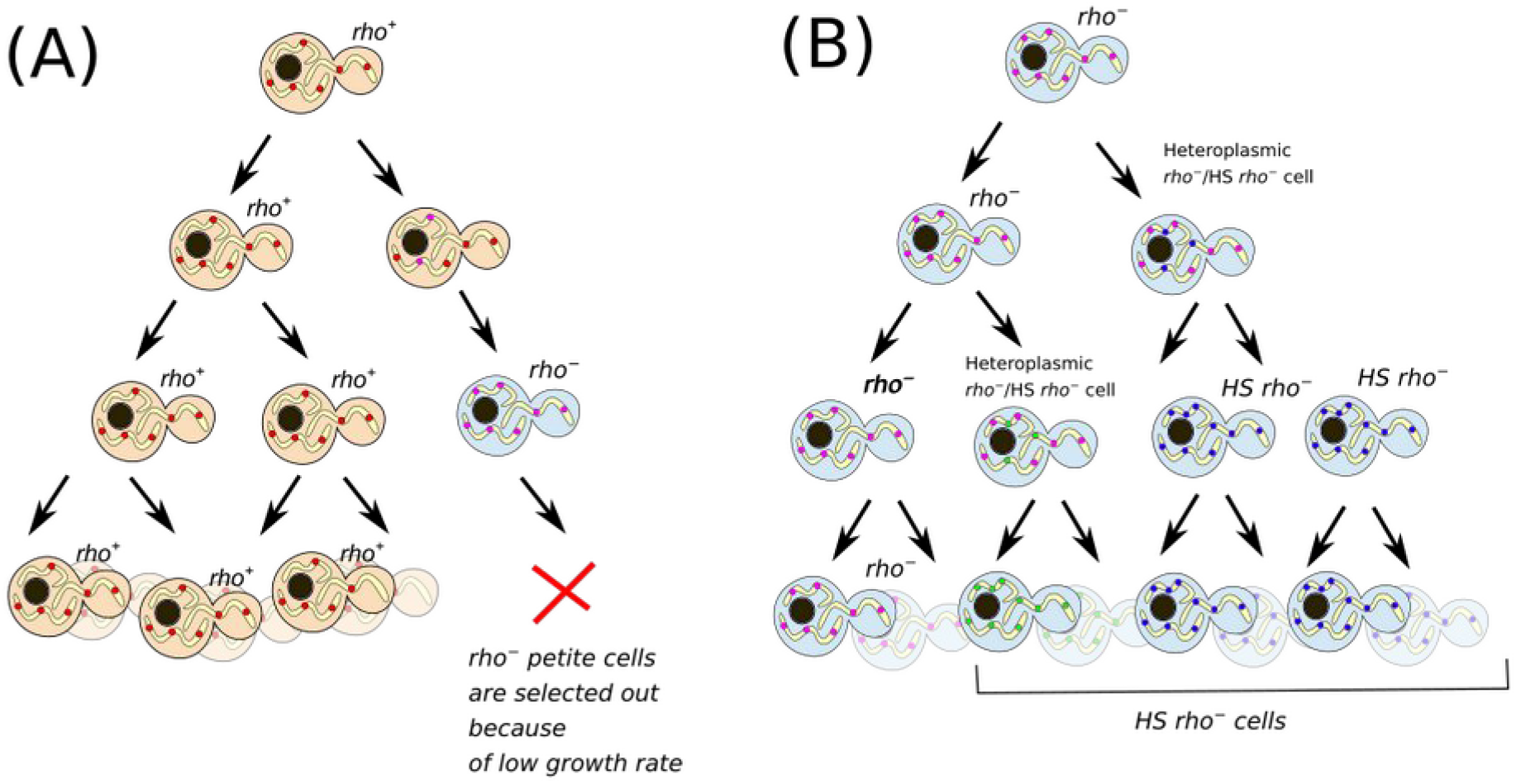
A schematic illustrating a hypothesis, where *rho*^−^ petite cells are continuously generated and selected out from the population of *rho*^*+*^ cells (A), while intracellular selection drives the fixation of highly suppressive (*HS*) *rho*^−^ variants within the *rho*^−^ petite cell population (B).

What could be the mechanism ensuring *rho*^−^ mtDNA intracellular fitness advantage in heteroplasmic cells? First, the advantage may be attributed to the reduced replication time of mtDNA molecules harbouring large deletions but retaining replication origins. Similar to PCR reactions, the amplification of shorter mtDNA molecules is more robust and proceeds more efficiently, especially under conditions of nucleotide deficiency. Indeed, the probability of *rho*^*+*^ mtDNA displacement appears to be negatively correlated with the size of the remaining mtDNA sequence [38] and positively correlated with the density of replication origins retained in a deletion variant of mtDNA [37]. Furthermore, the partial inhibition of dNTP synthesis by the overexpression of *SML1*, encoding the inhibitor of ribonucleotide reductase *RNR1*, strongly increased, and its deletion decreased the suppressivity [39]. Therefore, it might be that *rho*^−^ mtDNAs simultaneously cause problems with the biogenesis of essential DNA replication components (e.g. nucleotides and ATP) and gain a replication advantage because of these deficiencies.

Second, mtDNA may gain an advantage via preferential diffusion of mutant mtDNAs to the daughter cells. Upon cell division, yeast daughter cells receive a limited number of mtDNAs amplificated from a few randomly selected clones. This leads to a rapid loss of heteroplasmy by yeast cells because the progeny receives only one or another mitotype [40,41]. This mechanism is likely to rely on the limited diffusion of mtDNA molecules and mtDNA-encoded products along the mitochondrial network. The limited diffusion creates a linkage between mtDNA genotypes and phenotypes, which is considered to be essential for quality control of mtDNAs [42,43]. Cristae appears to be an essential factor preventing the diffusion of protein components of large protein complexes along mitochondria [44]. Accordingly, deletion of MIC (mitochondrial cristae organisation) genes prevent preferential inheritance of wild type mtDNA to a bud [45]. We speculate that *rho*^−^ mtDNAs showing intracellular advantage can avoid this sequestration, e.g. by the structural features of the spatial structure within the nucleoid. As a result, in contrast to wild type mitotypes, all daughter cells can receive a portion of mutant mtDNAs. Whatever the mechanism, our study suggests that the inheritance bias of *rho*^−^ mtDNAs is affected by variation in the intracellular fitness of the mutant mitotypes.

Biparental mtDNA inheritance repeatedly produces mitochondrial heteroplasmy, which creates the conditions for selfish mitochondrial genetic elements to spread in the fungal populations. Here, we provide rough estimates of the intracellular fitness ratio of mtDNA variants with deletions. Our data demonstrates that yeast mtDNA variants with large deletions can exhibit high intracellular fitness, challenging the fungal cells’ ability to maintain a functional mitochondrial genome containing all essential components for translation and oxidative phosphorylation.

## Methods

### Yeast strains, growth conditions and reagents

All strains used in this study are descendants from *W303-1A mat a* or *W303-1B mat alpha* stains used in our previous study [22]. Rich and complete synthetic media compositions were prepared according to Sherman [46] and manufacturer’s instructions (Sigma-Aldrich).

### Yeast DNA isolation & Quantitative PCR

To measure mtDNA/nDNA ratio, we took exponentially growing yeast cells and isolated total DNA using standard protocol which is based on the preparation of protoplasts and their lysis using SDS [47]. We carried out qPCR on the CFX96 Touch Real-Time PCR Detection System according to the manufacturer’s instructions. To quantify PCR products we used EvaGreen-ROX (Syntol) as the dsDNA probe. To detect mtDNA copy numbers in *rho*^−^ mutants, we designed three pairs of primers annealing to the different loci of the mtDNA genome (Table S3). As a reference to the nuclear DNA we used primers to the *ACT1* gene (Table S3).

### Library preparation and next generation sequencing of mitochondrial genomes

HS *rho*^−^ strain was sequenced using Illumina MiSeq platform and MiSeq Reagent Kit Version 3 (150 cycle). All other strains were sequenced using Illumina HiSeq platform, using HiSeq Flow Cell v3, TruSeq SBS Kit v3 (50-cycles) and TruSeq PE Cluster Kit v3 and on the Illumina Nextseq platform, using the Nextseq Mid Output Kit (150 cycle, paired-end sequencing). Libraries were prepared using the TruSeq RNA Library Prep Kit or NEBNext® Ultra™ II DNA Library Prep Kit for Illumina.

Obtained reads were trimmed with trimmomatic v. 0.32 with defaults parameters [48] and mapped with bowtie2 [49] to the reference S288C *Saccharomyces cerevisiae* genome (sacCer3, NCBI RefSeq assembly GCF_000146045.2).

### Quantification of mtDNA to nDNA ratio using Next Generation Sequence data

To quantify mtDNA copy numbers, we compared reads coverage to the mitochondrial and nuclear genomes; this approach has been previously used in other studies [23,50]. First, we mapped the sequence reads to the reference *S. cerevisiae* genomes using bowtie2 [49]. After this, we calculated nDNA average read coverage using Rsamtools (https://bioconductor.org/packages/Rsamtool). To negate the effect of rRNA repeat and telomeric regions which have higher read counts, we discarded nuclear genomic regions with extreme quantiles from the analysis. Thereafter, we calculated average mitochondrial genome read coverage. Given poor coverage of AT-rich genomic regions, to quantify mitochondrial genome read coverage we consider the coverage only the GC-rich regions of mitochondrial genome. Moreover, we calculated the coverage in the positions used for the quantitative PCR (see Figure S1). Given that *rho*^−^ strains retained random regions of the parental genome, we calculated the average coverage in all three regions, and then took the maximum of the obtained values. Relative read depth in different regions of mtDNA is presented in Table S2.

### Suppressivity assessment

Suppressivity was estimated as the proportion of *petite* diploid colonies in the total number of diploid progeny. Strains of two different mating types were grown to exponential phase in YPD and mixed in a 2 mL eppendorf tube or 96-well plate to the final OD = 0.1 of each strain (2 × 10^6^ cells/mL) in the total volume of 200 µL. Matings were performed at room temperature without mixing in rich media (YPD) if not indicated otherwise. After 20 h of mating cell suspensions were diluted with mQ water (10-1000 times depending on the mating efficiency for the given pair of strains) and plated on double-selective media (e.g. YNB-LEU-URA) containing 2% glycerol and 0.1% glucose as carbon sources. Such media allows selecting only diploid cells carrying both selective markers and furthermore distinguishing *petite* and *grande* diploid colonies. After 2-3 days we counted the number of *petite* and *grande* colonies and calculated the suppressivity.

### Stochastic model

To simulate genetic drift and dual level of selection in yeast we wrote a stochastic model using Julia language. The script is available here: (https://github.com/dknorre/YeastHeteroplasmy and illustrated in Figure S5. The algorithm is as follows: (1) A set of cells with specified levels of *rho*^−^ and *rho*^*+*^ molecules is generated. Each cell receives a random age parameter indicating the time spent after the previous division. (2) For each cell algorithm calculated the phenotype which was set as the function of *rho*^−^/*rho*^*+*^ mtDNA molecules ratio. In this study we assumed the threshold dependencies of the phenotype on the ratio of *rho*^*+*^/*rho*^−^ mtDNAs, although to improve robustness we tested different values of the threshold (Figure S6). In each simulation *petite* and *Grande* phenotypes were represented by duplication time, i.e. the number of minutes necessary to produce a new bud. The values were taken from the experiments in Figure 2 and set as 140 minutes for *Grande* phenotype and 210 minutes for *petite* phenotype. (3) Simulation increments yeast age for cell population. Each cell that reached its duplication time (according to the phenotype) divided and produced two cells with randomly distributed *rho*^−^ and *rho*^*+*^ mtDNAs. To do this we randomly sampled *rho*^*+*^ and rho^−^ mtDNAs into the new cells. Importantly, before the sampling we doubled the number of *rho*^*+*^ mtDNAs and multiplied the copies of *rho*^−^ mtDNAs copies by 2 * Intracellular fitness values. (4) After this, we returned to step two and repeated the simulation for 1440 minutes.

### Data analysis and visualisation

All data analysis and generation of figures were performed using R language version 4.4.0 and tidyverse libraries [51]. Circular map with mtDNA deletion variants were generated with circlize R package [52]

## Data Availability

All data are present within the manuscript and supplementary materials. Short-read sequencing results of *rho*^−^ strains are available in the SRA archive: BioProject PRJNA1039586. Julia language script for simulation of mtDNA drift in yeast clonal populations is available by the link: https://github.com/dknorre/YeastHeteroplasmy

## Authors contribution

DK proposed the project and acquired funding. NK and JK produced the mutant strains. NK isolated total yeast DNA, EG and ML prepared libraries and sequenced the yeast genomes. NK, SG, and DK analysed the NGS data. NK verified deletion positions using Sanger sequencing, performed qPCR experiments, suppressivity assays, and analysed growth rates. JK performed ethidium bromide sensitivity analysis. DK developed a stochastic model. NK and DK curated the data and prepared the illustrations. DK drafted the manuscript and supervised the research. All authors participated in discussing the results, writing and editing the text, and provided final approval of the version to be published

## Acknowledgements

We are grateful to Maria Kasyanova for the valuable help in confirming deletion sites in mtDNAs and Ekaterina Smirnova for conducting some of the crossing experiments.

