## Supplementary Information for "Inheritance bias of deletion-harbouring mtDNA in yeast: the role of copy number and intracellular selection"

Table S1. Deletion coordinates of *Rho⁻* strains. The coordinates were deduced from short-read sequencing assemblies and for some strains verified by PCR and Sanger sequencing.

| **Strain N** | **Coordinates of remaining mtDNA (start-end, b.p.)#** | **Primers used for deletion position verification/comments** | **Suppressivity (mean ± SD)** |
|---|---|---|---|
| *HS Rho⁻* | **31972-34167** | Isolated and characterised previously (Karavaeva et al. 2017) | 99.24 ± 0.42 |
| *Rho⁻ 2* | **79139-11106** | Forward: 5'-CCTGCGATTAAGGCATGATGA-3'<br>Reverse: 5'-GAATTTCGGTGATTGGAACC-3' | 47.61 ± 4.3 |
| *Rho⁻ 4* | **81404-35820** | Forward: 5'-CCTGCGATTAAGGCATGATGA-3'<br>Reverse 5'-ATGATAGATATCTGGGGTCC-3' | 50.38 ± 5.96 |
| *Rho⁻ 5* | 84230-34944 | Forward: 5'-GGATCAAACATTACCCGTTG-3'<br>Reverse: 5'-CATGACCCTAAAATGTTAACC-3'<br><br>PCR produced multiple products (250bp and 2500bp). Exact deletion coordinates were not verified and taken from illumina sequencing assembly. | 90.21 ± 5.14 |
| *Rho⁻ 6* | **79139-11106** | Forward: 5'-CCTGCGATTAAGGCATGATGA-3'<br>Reverse: 5'-GAATTTCGGTGATTGGAACC-3' | 68.41 ± 9.64 |
| *Rho⁻ 9* | 28455-6193 | | 68.76 ± 3.38 |
| *Rho⁻ 10* | 34762-56343 | | 76.64 ± 2.7 |

| | | | |
|---|---|---|---|
| *Rho⁻ 11* | 22690-35669 | | 88.1 ± 8.98 |
| *Rho⁻ 12* | 13391-35721 | | 89.17 ± 2.66 |
| *Rho⁻ 13* | 6205-52357 | | 44.95 ± 10.95 |
| *Rho⁻ 14* | **6987-52813** | Forward: 5'-GGAAATATAAAAACCGAAGG-3'<br>Reverse: 5'-CTGCTGGCACAAATATTAGTC-3' | 74.44 ± 15.79 |
| *Rho⁻ 15* | 30305-82529 | | 74.41 ± 6.18 |
| *Rho⁻ 18* | 29349-9065 | | 53.93 ± 12.15 |
| *Rho⁻ 19* | 71153-34184 | | 82.06 ± 7.38 |
| *Rho⁻ 20* | 73258-34273 | | 87.15 ± 5.55 |
| *Rho⁻ 21* | **35103-55705** | Forward: 5'-CTGCAATATCTTTTGCATTTG-3'<br>Reverse: 5'-GATGTCGTAACCATTAGACG-3' | 28.06 ± 9.5 |
| *Rho⁻ 22* | 70617-12320 | | 70.77 ± 0.68 |
| *Rho⁻ 45* | 53915-82908 | | 77.8 ± 5.55 |

| | | | |
|---|---|---|---|
| *Rho*$^{-}$ *Ia14* | 41776-66304 | | 43.85 ± 9.75 |
| *Rho*$^{-}$ *Ib28* | 28052-67035 | | 45.18 ± 17.2 |
| *Rho*$^{-}$ *IIa3* | 41659-4171 | | 44.92 ± 13.38 |
| *Rho*$^{-}$ *IIa10* | 8626-28624 | | 19.37 ± 8.06 |
| *Rho*$^{0}$ *Ib24* | n.d. appeared to be *rho*$^{0}$ | | 22.14 ± 16.01 |
| *Rho*$^{-}$ *IIc11* | 29409-4169 | | 63.11 ± 3.41 |

[#]coordinates according to the S288C reference genome annotation

Table S2. Quantification of mtDNA copy number from NGS data: depth of reads mapped to mtDNA normalised to depth of reads mapped to the nuclear genome. Trim Mean and median values were calculated for the entire mitochondrial genomes. In the rest of the columns, the average read depth was calculated in specified positions.

| **Strain** | Mean (trim = 0.1) | Median | Mean, primer set 1 position (31222-31305) | Mean, primer set 2 position (8002-8153) | Mean, primer set 3 position (48195-48296) | Mean, GC rich region (14100-21100) |
|---|---|---|---|---|---|---|
| *HS Rho⁻* | 40.13 | 0.04 | 679.69 | NA | NA | NA |
| *Rho⁻ 2* | 10.16 | 9.96 | NA | 24.10 | NA | 0.03 |
| *Rho⁻ 4* | 9.94 | 5.08 | 87.76 | 6.46 | NA | 30.82 |
| *Rho⁻ 5* | 2.00 | 1.32 | 183.35 | 0.97 | NA | 2.44 |
| *Rho⁻ 6* | 11.55 | 6.11 | 46.19 | 0.08 | 33.04 | 0.47 |
| *Rho⁻ 9* | 10.08 | 8.34 | 41.38 | NA | 14.70 | 0.03 |
| *Rho⁻ 10* | 7.59 | 6.50 | NA | NA | 12.72 | 0.01 |
| *Rho⁻ 11* | 16.29 | 0.75 | 235.83 | NA | NA | 0.07 |
| *Rho⁻ 12* | 2.79 | 0.89 | 68.96 | NA | NA | 0.83 |
| *Rho⁻ 13* | 8.46 | 4.51 | 10.32 | 3.72 | 3.85 | 24.28 |
| *Rho⁻ 14* | 4.81 | 2.95 | 10.97 | 35.33 | 4.56 | 2.57 |
| *Rho⁻ 15* | 5.45 | 3.92 | 28.55 | 0.04 | 5.32 | 0.04 |
| *Rho⁻ 18* | 10.63 | 8.06 | 27.52 | 3.62 | 21.43 | 0.14 |
| *Rho⁻ 19* | 6.60 | 3.52 | 821.52 | 0.97 | NA | 13.10 |

| | | | | | | |
|---|---|---|---|---|---|---|
| *Rho$^{-}$ 20* | 2.36 | 1.22 | 187.83 | 0.53 | NA | 4.79 |
| *Rho$^{-}$ 21* | 7.76 | 5.41 | 0.10 | NA | 13.11 | 0.04 |
| *Rho$^{-}$ 22* | 46.62 | 45.09 | 0.21 | 130.47 | 0.40 | 0.21 |
| *Rho$^{-}$ 45* | 20.61 | 19.34 | 0.50 | NA | NA | 0.27 |
| *Rho$^{-}$* Ia14 | 7.66 | 6.90 | NA | NA | 21.45 | NA |
| *Rho$^{-}$* Ib28 | 10.63 | 9.98 | 9.85 | NA | 19.35 | NA |
| *Rho$^{-}$* IIa3 | 10.77 | 9.25 | 28.64 | NA | NA | 25.60 |
| *Rho$^{-}$* IIa10 | 12.15 | 10.12 | NA | NA | 25.72 | NA |
| *Rho$^{-}$ IIc11* | 59.13 | 53.35 | NA | 0.06 | 55.06 | NA |
| ***Rho$^{+}$ strain*** | 4.37 | 3.00 | 6.56 | 4.51 | 4.81 | 15.45 |
| *rho$^{0}$* strain | 0.19 | 0.19 | 0.23 | NA | NA | 0.20 |
| *rho$^{0}$* strain -2 | 0.08 | 0.07 | NA | NA | NA | NA |

Table S3. Primers, used for quantitative PCR of mtDNA to nDNA ratio

| Name | Sequence | Product, b.p. | Annealing temperature, |
|---|---|---|---|
| qPCR region 1 | Forward: att-cca-cct-tca-gcg-tag-t<br>Reverse: ggt-tcg-gtc-ctc-cct-tac | 83 | 60°C |
| qPCR region 2 | Forward: ttc-gca-cta-atc-act-cat-cac<br>Reverse: ccc-tac-ggt-aac-tgt-att-tca-ac | 152 | 60°C |
| qPCR region 3 | Forward: gta-tta-tta-cgg-atg-atg-tag-gat<br>Reverse: aag-gat-ggt-tga-ctg-agt | 98 | 53°C |
| ACT1 | Forward: tcc-cag-gta-ttg-ccg-aaa-gaa-tgc<br>Reverse: gcc-aag-ata-gaa-cca-cca-atc-cag-ac | 124 | 58°C |

Table S4. Quantification of mitochondrial DNA (mtDNA) copy number using qPCR analysis. The mtDNA copy number was determined by the ratio of mitochondrial to nuclear genome markers, calculated as -logCt. The amplified regions 1 and 3 used for mtDNA quantification are illustrated in Figure S1.

| **Strain** | mtDNA/ nDNA | mtDNA/nDNA (region 1), mean | mtDNA/nDNA (region 1), SD | mtDNA/nDNA (region 3), mean | mtDNA/nDNA (region 3), SD | mtDNA Rho- normalised to WT |
|---|---|---|---|---|---|---|
| *HS Rho⁻* | 686 | 686 | 216 | NA | NA | 28.3 |
| *Rho⁻ 4* | 206 | 206 | 84 | NA | NA | 8.5 |
| *Rho⁻ 5* | 357 | 357 | 99 | NA | NA | 14.7 |
| *Rho⁻ 6* | 62 | 70 | 45 | 56 | 16 | 2.6 |
| *Rho⁻ 9* | 55 | 63 | 34 | 48 | 24 | 2.3 |
| *Rho⁻ 10* | 107 | NA | NA | 108 | 25 | 4.4 |
| *Rho⁻ 11* | 341 | 341 | 129 | NA | NA | 14.1 |
| *Rho⁻ 12* | 197 | 197 | 77 | NA | NA | 8.1 |
| *Rho⁻ 13* | 50 | 49 | 17.5 | 52 | 26 | 2.1 |
| *Rho⁻ 14* | 34 | 48 | 16.4 | 24.5 | 10 | 1.4 |
| *Rho⁻ 15* | 68 | 102 | 58 | 46 | 11 | 2.8 |
| *Rho⁻ 18* | 42 | 41 | 7.9 | 43 | 12 | 1.7 |
| *Rho⁻ 19* | 412 | 412 | 106 | NA | NA | 17.0 |

| | | | | | | |
|---|---|---|---|---|---|---|
| *Rho⁻ 20* | 233 | 232 | 57 | NA | NA | 9.6 |
| *Rho⁻ 21* | 124 | NA | NA | 124 | 59 | 5.1 |
| ***Rho⁺ strain*** | 24 | 22 | 4.4 | 26.6 | 9.4 | 1 |

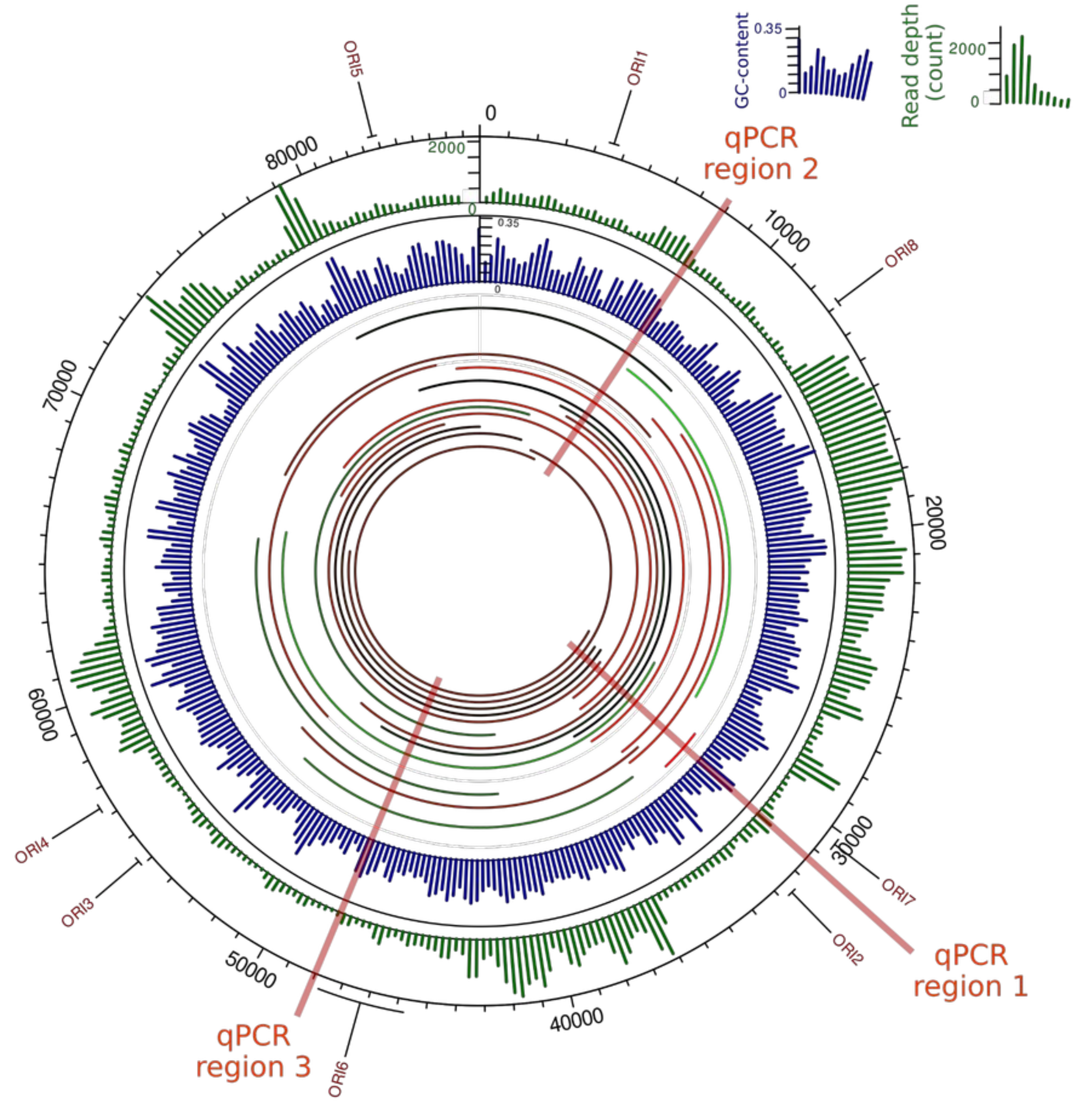


Figure S1. Relative position of the regions used for mtDNA quantitative PCR and the regions retained in *rho*⁻ mitochondrial genomes.

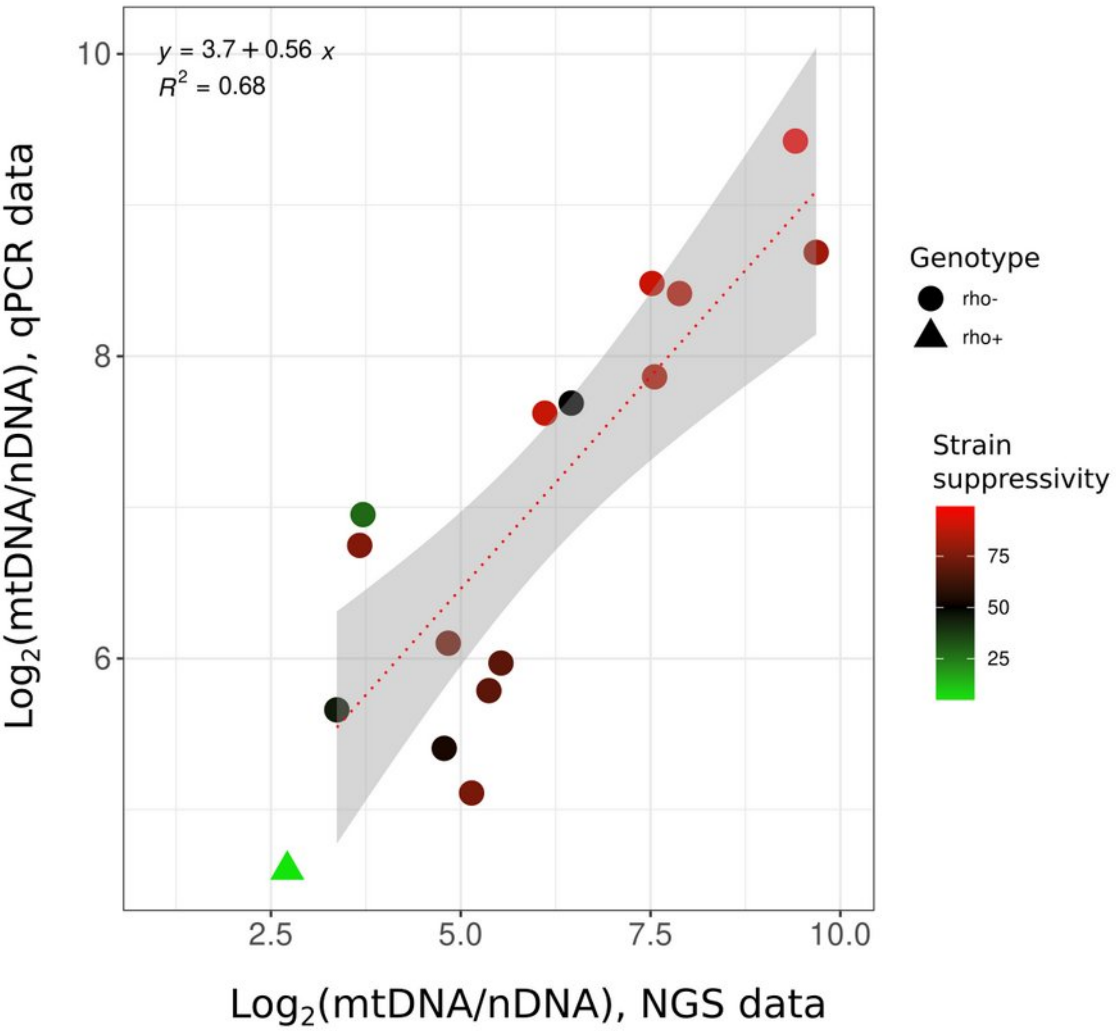


Figure S2. Concordance between mean qPCR estimates of mtDNA copy numbers and NGS-based estimates. Kendall's rank correlation tau = 0.683, p-value = 8.266e-05.

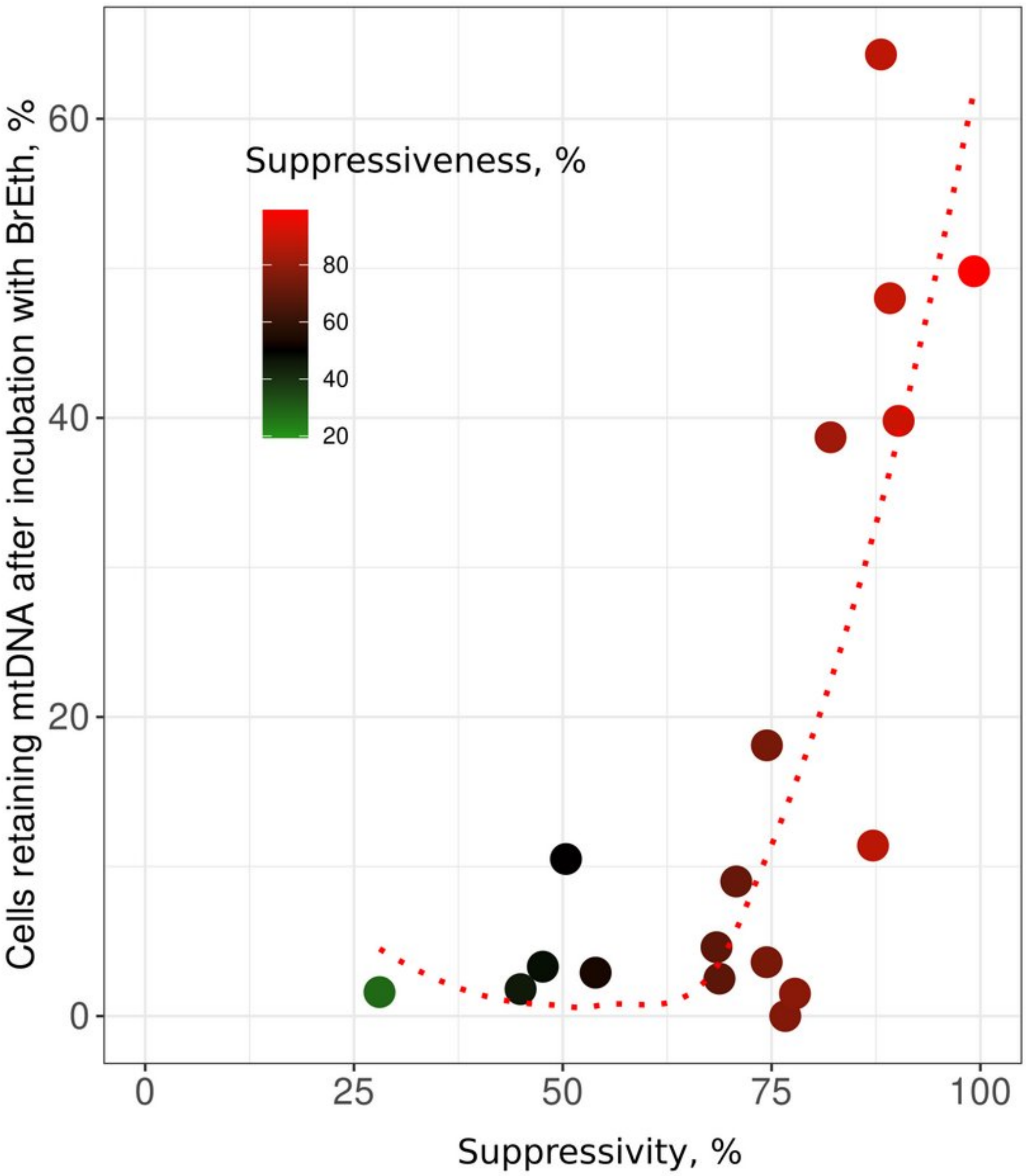


Figure S3. The ability of *rho⁻* yeast cell to retain mtDNA upon growth with the DNA-intercalating agent ethidium bromide correlates with rho-strain suppressivity (Kendall's rank correlation tau = 0.56, p-value = $8*10^{-4}$)

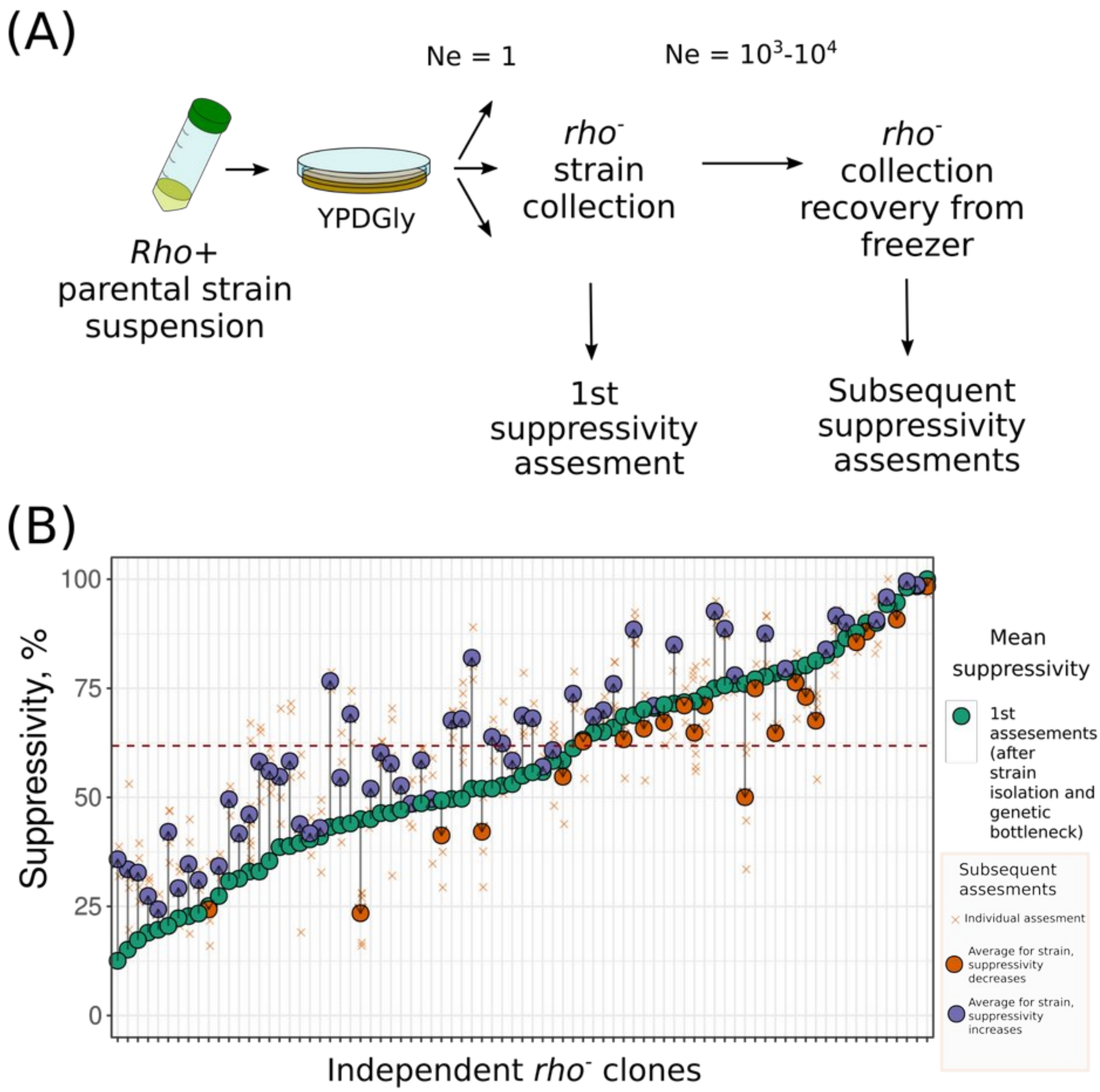


Figure S4. Suppressivity increases within yeast subclones. (A) Schematic showing the experimental setup for measurement of suppressivity drift over generations; (B) Change in suppressivity between first (green circles) and subsequent (orange and blue circles) assessments (n = 76). The orange vs blue color of the circle illustrates the direction of change in subsequent experiments compared to the first one, with orange indicating a decrease and blue indicating an increase of suppressivity over generations. X labels the results of individual experiments.

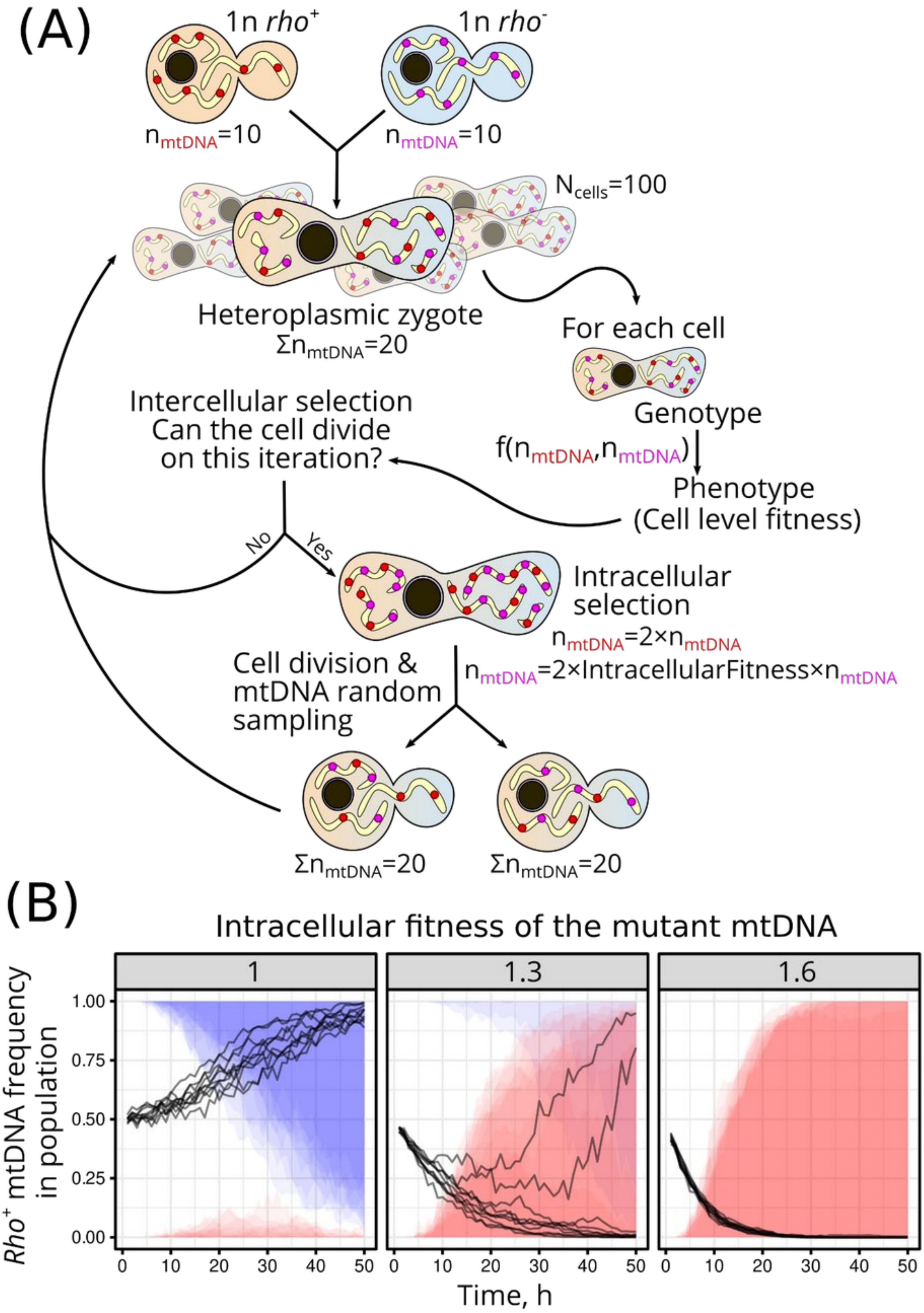
(A)
1n rho+
1n rho-
nmtDNA=10
nmtDNA=10
Ncells=100
Heteroplasmic zygote
ΣnmtDNA=20
For each cell
Genotype
f(nmtDNA,nmtDNA)
Phenotype
(Cell level fitness)
Intercellular selection
Can the cell divide
on this iteration?
No
Yes
Intracellular
selection
nmtDNA=2×nmtDNA
nmtDNA=2×IntracellularFitness×nmtDNA
Cell division &
mtDNA random
sampling
ΣnmtDNA=20
ΣnmtDNA=20
(B)
Intracellular fitness of the mutant mtDNA
1
1.3
1.6
Rho+ mtDNA frequency
in population
1.00
0.75
0.50
0.25
0.00
0
10
20
30
40
50
Time, h

Figure S5. MtDNA selection in heteroplasmic yeast cells: a simulation guided by experimental data. (A) Simulation algorithm, see details in material and methods section. (B) The simulation predicts the proportion of cells containing only *rho*$^{+}$ mtDNA (represented by blue area) and those with only *rho*$^{-}$ mtDNA (represented by red area). Each plot represents an overlay of the results of ten simulations, to overlay blue and red areas that illustrate homoplasmic cells we set their opacity to 0.1. Not coloured areas correspond to the cell retaining heteroplasmy. nd *rho*$^{-}$ cells. Three plots represent simulation results with different intracellular fitness levels of rho- mtDNA (1, 1.3, and 1.6). The relative cell-level fitness of *rho*$^{-}$ cells was set at 0.75 for this simulation. The pathogenicity threshold was set at 0.5, indicating that a *rho*$^{-}$ cell-level phenotype only manifests when more than half of the mtDNA in a cell is rho-.

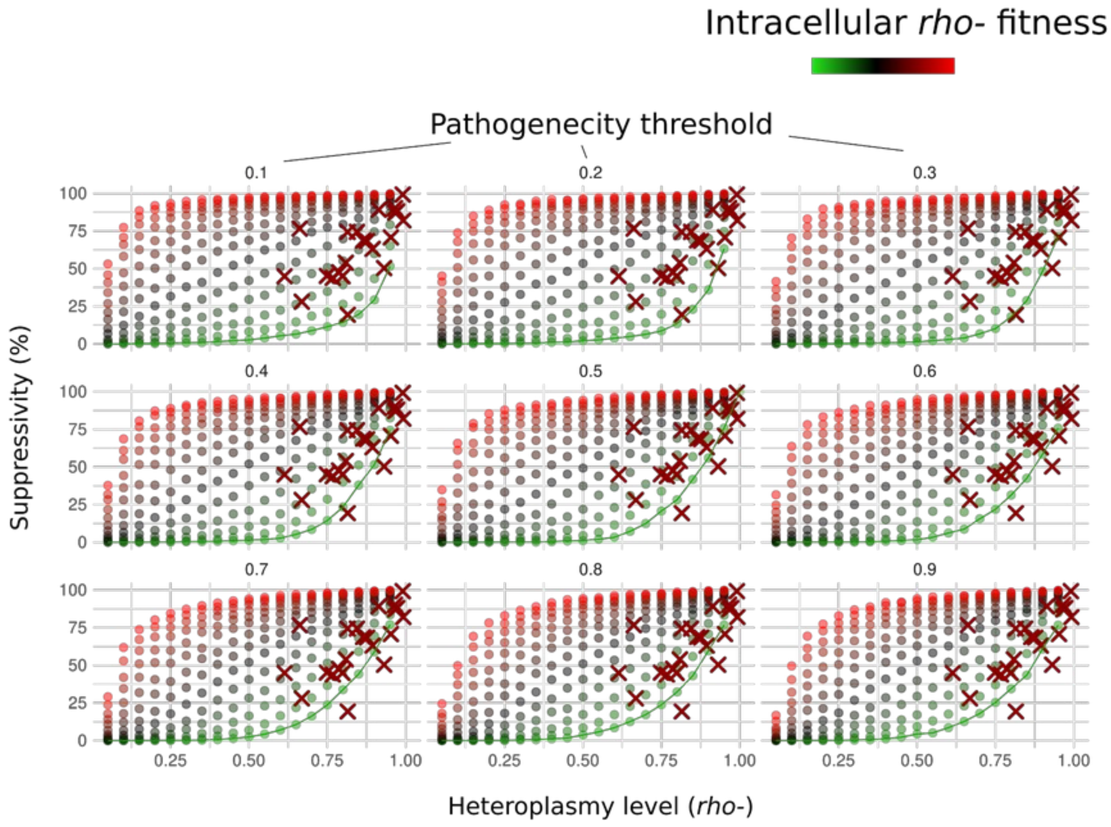

Figure S6. Variation of pathogenicity threshold in simulation of suppressivity. Simulation data and experimental data plotted in the coordinates of starting heteroplasmy level ~ suppressivity (same as in the Figure 4B); Points represent simulated data, crosses represent experimental data points for the 22 *rho*⁻ strains; a line connects the points obtained with the simulation with no replication advantage of *rho*⁻ mtDNA (Intracellular *rho*⁻ mtDNA fitness equal to 1.0)

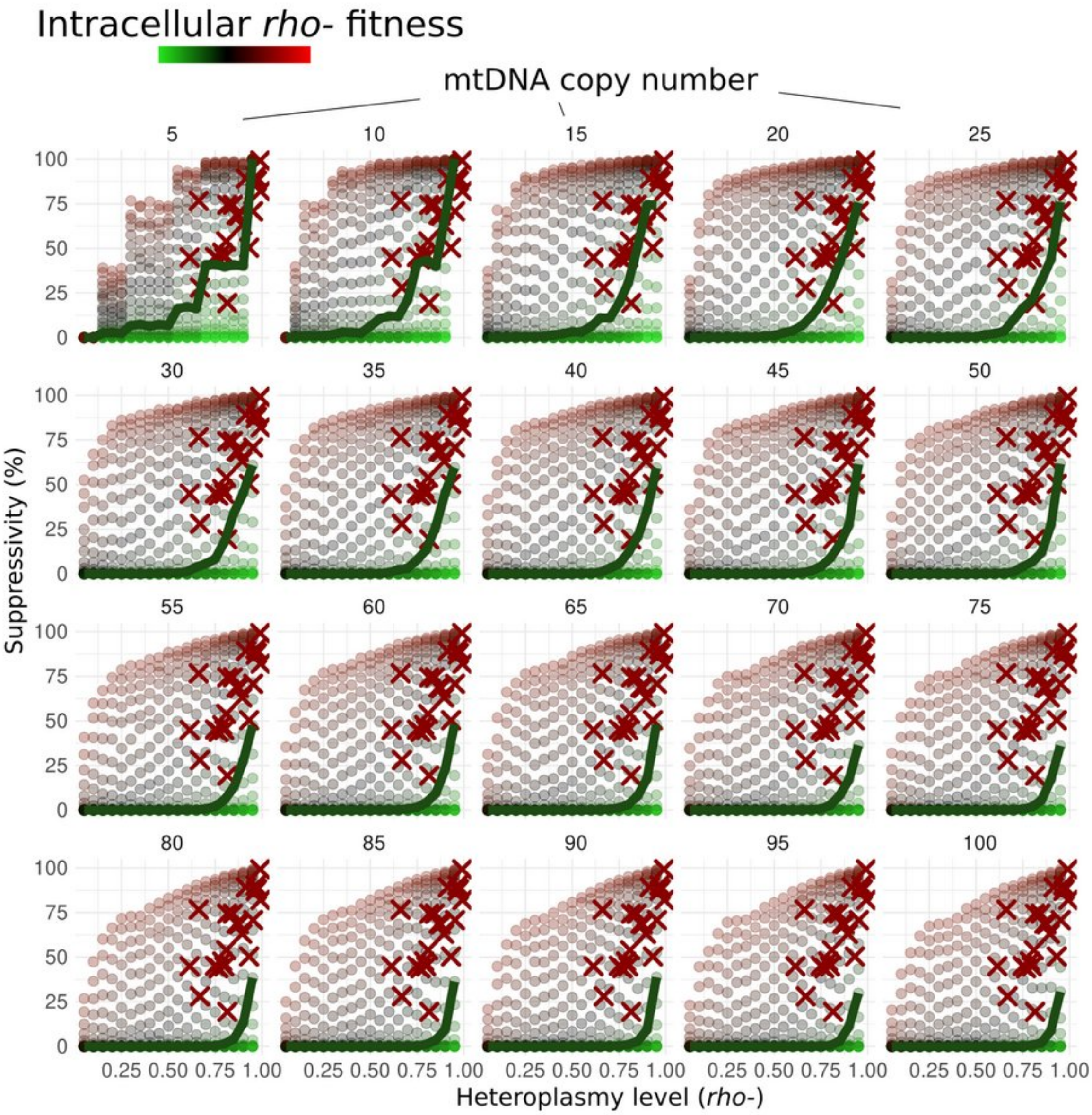

Figure S7. Variation of mtDNA copy number ($n_{mtDNA}$) in simulation of suppressivity. Simulation data and experimental data plotted in the coordinates of starting heteroplasmy level ~ suppressivity (same as in the Figure 3B); Points represent simulated data, crosses represent experimental data points for the 22 *rho*⁻ strains; a line connects the points obtained with the simulation with no replication advantage of *rho*⁻ mtDNA (Intracellular *rho*⁻ mtDNA fitness equal to 1.0)

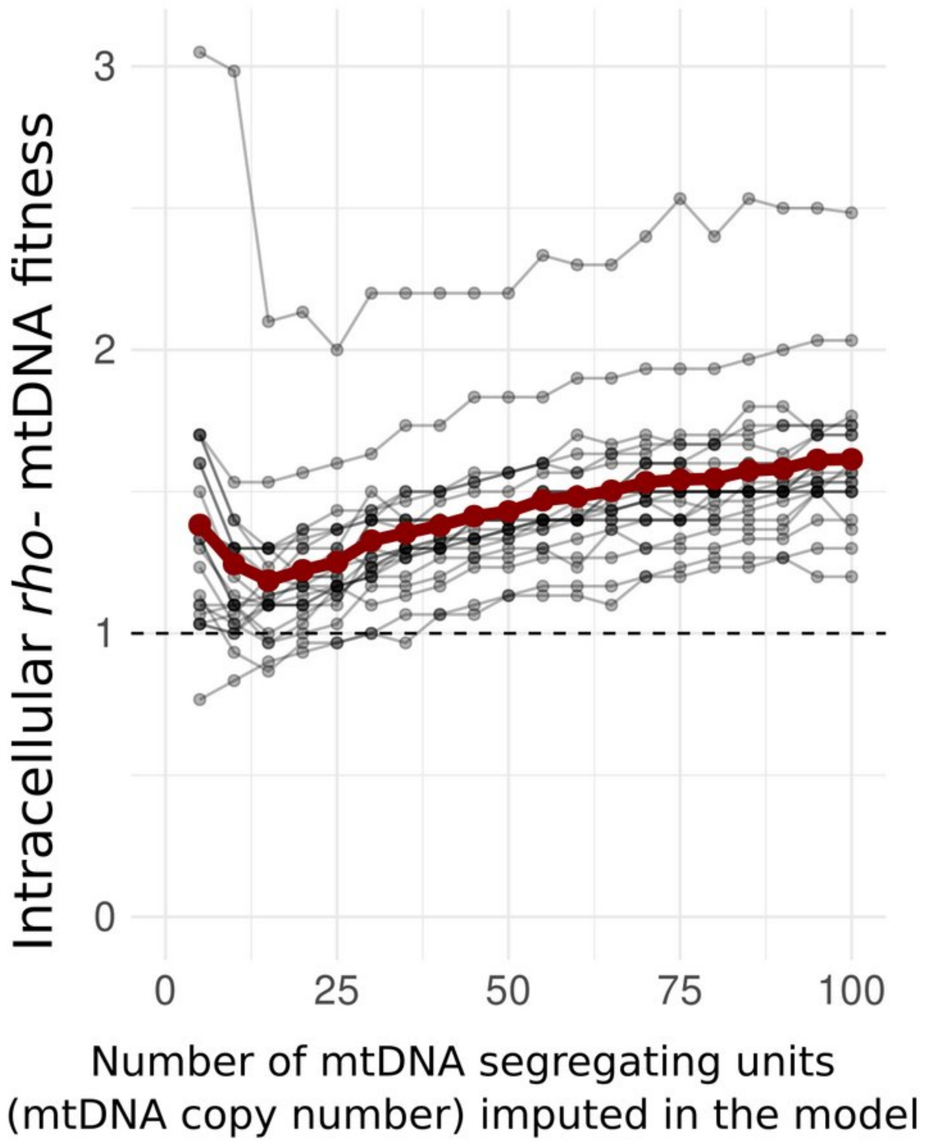

Figure S8. Relative intracellular fitness of *rho*$^{-}$ mtDNA variants calculated from the k-nearest neighbours for simulations with different numbers of mtDNA copy number (number of mtDNA segregating units). Individual lines illustrate the predictions for individual *rho*$^{-}$ strains, the red bold line shows the average value.
